# Eastern Baltic cod larvae in a salinity gradient: choice of salinity and the role of neutral buoyancy

**DOI:** 10.1101/2025.09.18.677118

**Authors:** Maddi Garate-Olaizola, Johanna Fröjd, Jane W. Behrens, Sune Riis Sørensen, María Cortázar-Chinarro, Anssi Laurila

## Abstract

Fish rely on multiple sensory systems to navigate in the aquatic environment, yet these mechanisms are often scarcely known during their early life history stages. Salinity is an important environmental parameter affecting the performance of fish larvae, and recent studies have shown that some larval fish use it as an environmental cue to position, orientate and navigate. Further, fish larvae are often visual feeders, and light has been suggested to be a crucial environmental cue for visual orientation and feeding. In this study, we investigate whether first-feeding larvae of eastern Baltic cod (*Gadus morhua callarias*) are driven by light and salinity. We examine this in the context of their natural vertical distribution and movement; from deeper, darker and more saline (13-17 psu) water layers towards the more light-exposed and less saline (7 psu) surface water. Further, we investigate the relationship between this movement and the neutral buoyancy measured both in active and passive larvae. We found that the larvae remain in a saline environment close to the salinity which supports their neutral buoyancy. They show negative phototaxis, suggesting an active avoidance of the lighter and less saline upper water layers. These results improve our understanding of the ecophysiology and behaviour of eastern Baltic cod larvae during the first-feeding stage and serve as an important input for restocking efforts of eastern Baltic cod and potential future production of hatchery rearing of eastern Baltic cod.

**Highlights:**

- First-feeding larvae had no preference for a low-salinity environment.
- Vertical distribution of eastern Baltic cod larvae was mainly driven by negative phototaxis.
- Larval neutral buoyancy played a larger role in their behaviour than light.

## Introduction

Fish use multiple cues and sensory systems to orientate and navigate in the aquatic environment (Facey et al., 2022a), however, most research deals with sensory mechanisms in the adult stage and relatively few studies focus on the early life history stages. Earlier studies show that fish larvae can use multiple sensory systems such as olfaction, vision, hearing and even magnetic detection for orientation and to find suitable settlement habitats (Arvedlund and Kavanagh, 2009; Baptista et al., 2020; Berenshtein et al., 2022; Leis, 2006). Some fish larvae and juveniles navigate towards estuarine nursery areas based on multiple environmental cues like odours, temperature and salinity gradients (Boehlert and Mundy, 1988).

In marine habitats, salinity is an important environmental parameter for fish, which detect it through special receptors (Nearling et al., 2002). Salinity changes will challenge osmoregulation, and here fish larvae are particularly sensitive given the early developmental stage with less developed osmoregulatory systems, a large surface area to body mass ratio and limited movement ability. This makes coping with salinity changes harder and can lead to rapid dehydration or overhydration depending on the salinity of their environment (Govoni and Forward Jr, 2008). Large shifts in salinity can occur in marine environments, especially in the presence of a vertical halocline (Bashevkin et al., 2020). For example in the eastern Baltic, water with lower salinity is restricted to the surface due to differences in water density, while higher salinity water is found in the deep water (Snoeijs-Leijonmalm and Andrén, 2017). It has been suggested that teleosts use salinity gradients as a navigational cue to avoid adverse environmental conditions and osmotic stress, which can impact growth and even survival in both marine and freshwater fishes (Arvedlund and Kavanagh, 2009; Bœuf and Payan, 2001, p. 20; Herrera et al., 2021; Serrano et al., 2010). Selection for suitable habitats based on salinity has been found in larval apogonids and pleuronectids (Atema et al., 2002; Dando, 1984), and a recent study showed that larval zebra fish detect salinity in the water through the olfactory system, triggering a behavioural response towards favourable concentrations (Herrera et al., 2021). Despite that functional mechanisms behind salinity detection in fish larvae are largely unknown, it seems evident that fish larvae can detect and respond to salinity gradients (Burke et al., 1995).

The ecology of Atlantic cod larvae (*Gadus morhua*), particularly the foraging and locomotory behaviour, is poorly understood despite the high commercial importance of the species. The eastern stock of the Baltic cod *Gadus morhua callarias*, is a phenotypically and genetically isolated subspecies of the Atlantic cod (Barth J. M. I. et al., 2019; Berg et al., 2015; Helmerson et al., 2023; Kraus, 2002; Nissling and Westin, 1997). This stock has been heavily exploited by fisheries for several decades, which, together with unfavourable environmental conditions, parasite infestation and reduced food availability led to population collapse in the late 1900s and the total closure of both commercial and recreational fisheries in 2019 (Eero et al., 2023, 2015). Historically suitable spawning grounds of eastern Baltic cod covered several deep basins in the Baltic Sea, however, many of these were lost due to intensified hypoxic conditions in the Baltic deep waters affecting multiple spawning grounds and restricting its reproduction to the Bornholm Basin, located in the southern part of the Baltic Sea (Almroth-Rosell et al., 2021; Casini et al., 2016; Eero et al., 2023; Hinrichsen et al., 2016).

Yolk-sac larvae of Atlantic cod hatch from pelagic eggs (Nissling and Westin, 1991a), which remain suspended in the water column throughout embryonic development due to their buoyancy. Egg buoyancy is a consequence of the difference in osmolarity between the egg and the surrounding water, and it is determined by maternal fluids (absorbed by the oocyte during oocyte maturation in the ovary) and facilitated by yolk lipids and oil droplets (Fabra et al., 2005; Govoni and Forward Jr, 2008). These provide the eggs with specific density (Govoni and Forward Jr, 2008; Nissling and Westin, 1991a). At the same time, the density of the environmental water is influenced by its salinity (among others), becoming denser as the salinity increases and vice versa (Dougherty, 2001; Govoni and Forward Jr, 2008). The relation between the specific density of the egg with the density of the environmental water will determine whether the eggs float or sink (Nissling and Westin, 1991a). When egg specific density matches the density of the surrounding water at a particular environmental salinity, the eggs reach “*neutral buoyancy*” (Govoni and Forward Jr, 2008; Nissling and Westin, 1991a). In this state, eggs are neutrally buoyant, remaining suspended in the water column, and the environmental salinity (psu) at which this happens quantifies their neutral buoyancy. (Nissling et al., 2017, 1994a; Nissling and Westin, 1991a; Saborido-Rey et al., 2003).

The Baltic Sea is a brackish environment, with a permanent halocline at 50-60 m in the Bornholm basin, separating a low saline (6-8 psu) water mass above with a higher saline water mass (13-17 psu) below (Snoeijs-Leijonmalm and Andrén, 2017). Eastern Baltic cod eggs become buoyant at 14-15 psu (Nissling and Vallin, 1996), which delimits their depth of development to 60-80m, below the halocline and in a dark, more saline environment (Nissling and Westin, 1991a; Snoeijs-Leijonmalm and Andrén, 2017). Here, the temperature is 4-6 °C (Snoeijs-Leijonmalm and Andrén, 2017), and the yolk-sac larvae hatch approximately 12-13 days after fertilisation, and active swimming starts 4-6 days after hatching (Fossum, 1986; Hall et al., 2004). Concurrent with this, eyes, jaw, and a functional stomach develop, and exogenous feeding initiates (Ellertsen et al., 1980; Fossum, 1986; Hall et al., 2004; Støttrup et al., 2008).

The first-feeding cod larvae are 4-5 mm (Grønkjær & Wieland, 1997; Huwer, 2009) and rely on small prey, preferring copepod nauplii but also consuming dinoflagellates and phytoplankton (Ellertsen et al., 1980; Haldorson et al., 1993; Hall et al., 2004; Jakobsen et al., 2020; Marak, 1960; Van der Meeren and Naess, 1993). This prey is found in the photic zone above the halocline at 20-40m depth and shows dial vertical movement (Renz J. and Hirche H-J., 2006; Snoeijs-Leijonmalm and Andrén, 2017). Like most marine fish larvae, larval cod are visual feeders, relying on light to find food (Ellertsen et al., 1980; Hall et al., 2004) and are limited to relatively short distances to detect it (Colton and Hurst, 2010). Thereby, first-feeding larval eastern Baltic cod are expected to cross the halocline and swim upwards tens of meters in the water column to match their prey and to initiate feeding (Grønkjær et al., 1997; Grønkjær and Wieland, 1997), moving from a dark environment with higher salinity (13-17 psu) towards a lighter environment with lower salinity (6-8 psu). These assumptions are partially in line with available field studies (Grønkjær et al., 1997; Grønkjær and Wieland, 1997; Huwer, 2009). Despite the importance of environmental cues in larval cod navigation, little attention has been given to understanding which cues they rely on during this vertical migration. For instance, salinity as a navigational cue has not been investigated in larval eastern Baltic cod, nor has the interaction with light.

Atlantic cod larvae exhibit neutral buoyancy similar to that of eggs immediately after hatching; however, this buoyancy decreases as larvae grow, since it is influenced by body size (Ellertsen et al., 1980; Fossum, 1986; Govoni and Forward Jr, 2008; Saborido-Rey et al., 2003). A similar pattern is known in eastern Baltic cod (Støttrup et al., 2008). Neutral buoyancy of larval eastern Baltic cod is 14-17 psu after hatching, similar to the eggs and setting them below the halocline (Nissling and Vallin, 1996). As larvae grow, they become less buoyant (Ellertsen et al., 1980; Saborido-Rey et al., 2003). The movement of the larvae upwards is hence energetically costly but needed in order to reach sufficient food abundance. The active transition from salinity close to their neutral buoyancy towards the hypo-osmotic environment means an active process requiring energy. Due to the strong contrast in the environment above and under the halocline, salinity and their neutral buoyancy may influence the behaviour of cod larvae in the Baltic Sea. The link between larval cod behaviour, their neutral buoyancy and light has received little attention and needs to be studied in more detail.

This study used an experimental approach to investigate the movement of eastern Baltic cod larvae in an environment resembling the vertical salinity and light profile in the Baltic Proper (the main basin of the Baltic Sea). We hypothesized that cod larvae respond to salinity and light, thereby influencing their vertical positioning and movement patterns along these gradients. Since zooplankton is typically concentrated near the surface, where salinity is lower and light availability is higher, we hypothesize that cod larvae preferentially inhabit low-salinity, well-illuminated environments. We further investigated how the neutral buoyancy of eastern Baltic cod larvae is linked to these behaviours as well as to larval body size.

## Material and methods

### Broodstock acquisition and spawning

Adult eastern Baltic cod (n = 93) were caught by trawl in ICES subdivisions 25, 26 and 28 on four occasions between February 2021 and May 2022. The fish were brought to the research station Ar in northern Gotland, where they were distributed in two circular tanks with a water volume of 14 m^3^ (3.5 m in diameter with a water depth of 1.46 m) supplied with air-saturated recirculated water (7 °C, 17 psu). Each tank was connected to a water treatment unit made up of a drum filter, UV filter, biofilter, trickle-filter, protein skimmer and heat exchanger for temperature regulation. Ozone was automatically dozed in the protein skimmer if the ORP values were below 290 mV. Natural seawater was obtained from 35 m depth and filtered by a 0.5 μm particle filter and UV filter. The salinity was adjusted from 7 psu to 17 psu by using artificial sea salt (Hoss, fish salt, Karlslunde, Denmark). The light regime was set to follow the natural light regime in Visby, Gotland (57°37′44″N 18°18′26″E) with Philips aquaculture light system 125W, including simulation of dawn and dusk. During the experimental period (June-July 2023), the fish were fed *ad libitum* every second day with shrimp. Fish started to spawn in late April and were allowed to spawn freely (see Table S1 for an overview of parental fish). Eggs were collected once daily (each collection representing one temporal batch) during the spawning season using floating egg-collecting devices (Cortney Ohs et al., 2019).

### Egg incubation and hatching

Each batch, consisting of thousands of eggs of several females not older than 24 h post-fertilization, was transferred to a single 80L incubator (7 °C, 17 psu). The incubation system was made as a separate RAS unit with a water treatment system formed by filter socks, biofilter, protein skimmer and UV filter. Each incubator was stocked with up to 250 ml of buoyant eggs. A gentle aeration and water were supplied at the bottom of each incubator. The water exited via a banjofilter with a mesh size of 250 mμ and led back to the water treatment unit. Once a day, aeration and water were stopped, and dead eggs were purged from the bottom drain. The experiment was carried out using eggs from five temporally separated batches collected between the 4^th^ and 26^th^ of June 2023. Eggs were incubated until hatching, with 50% hatching estimated to occur on day 13. The larvae were reared for four days post-hatching, reaching developmental stages 4–5 (Ellertsen et al., 1980; Fossum, 1986) where they possess a developed functional jaw and are ready to start exogenous feeding (Ellertsen et al., 1980; Fossum, 1986; Hall et al., 2004). Different sets of 120 larvae belonging to the same batches were used in the behavioural and neutral buoyancy assessment.

### Experimental setup

The day before the experiment, experimental seawater of salinities 9, 11, 13, 15 and 17 psu was prepared by adjusting filtered (0.5 μm particle filter, Spectrum, England and Wales, UK) natural brackish seawater from the Baltic with artificial sea salt (Hoss, fish salt, Karlslunde, Denmark). Salinity and temperature were measured with an electronic conductivity meter (WTW Multi 3410 with TetraCon925 ids sensor, WTW Wissenschaftlich-Technische Werkstätten GmbH, Weilheim, Germany). Oxygen content was measured with 0.5 % accuracy using an oxygen probe (WTW FDO® 925 Optical IDS dissolved oxygen sensor) connected to the conductivity meter.

### Salinity choice experiment

The experiment was performed in a cold room at 7 °C. Three 300 ml glass cylinders (height 32.5cm, diameter 4.0 cm) were used for the salinity choice experiment: one cylinder with a salinity gradient from 7 psu on the top to 17 psu on the bottom and two cylinders having a uniform salinity, i.e. no gradient. The salinities in the latter two treatments were the upper and lower salinity limits (7 and 17 psu) (Figure 1). All three cylinders were divided into six vertical sections (Figure 1, see also below for the preparation of the salinity gradient). These sections were based on the different salinity layers in the salinity gradient and used to delimit the vertical space in the three cylinders so that the size of the sections in the control cylinders was identical to that in the gradient cylinder (Figure 1). The control cylinders (7 and 17 psu, respectively) acted as a reference for the position of the larvae in the gradient cylinder, the number of larvae in each section reflecting the spatial behaviour in the absence of the salinity gradient. The three cylinders were tested simultaneously, with a single trained observer who was blind to the salinity treatment in each cylinder. The cylinder was filled with up to 50 ml of 7 psu water from the bottom. Thereafter, 50 ml of 9 psu water was added very gently from the bottom to the cylinder, leaving the less saline water on the top. This process was repeated with salinities 11, 13, 15 and 17 psu. The process resulted in 300ml of water, divided into six salinity sections 30-55mm deep in the cylinder. To confirm and delimit the salinity gradient sections, five glass beads with particular colour patterns and specific neutral buoyancy (∼ 6 mm Diam; Martin Instrument Co. Welwyn Garden City AL7 4BG, Herts., England, UK) of 7.7, 9.7, 11.6, 14.5 or 16.6 psu were added to the gradient cylinder. As each glass bead positioned itself between two salinity sections, this could be used to delimit and mark the salinity sections, which was then used to align the placement of the larvae in the cylinder. The gradient was assessed post-experimentally to avoid any influence of the glass beads on larval behaviour.

**Figure 1:**
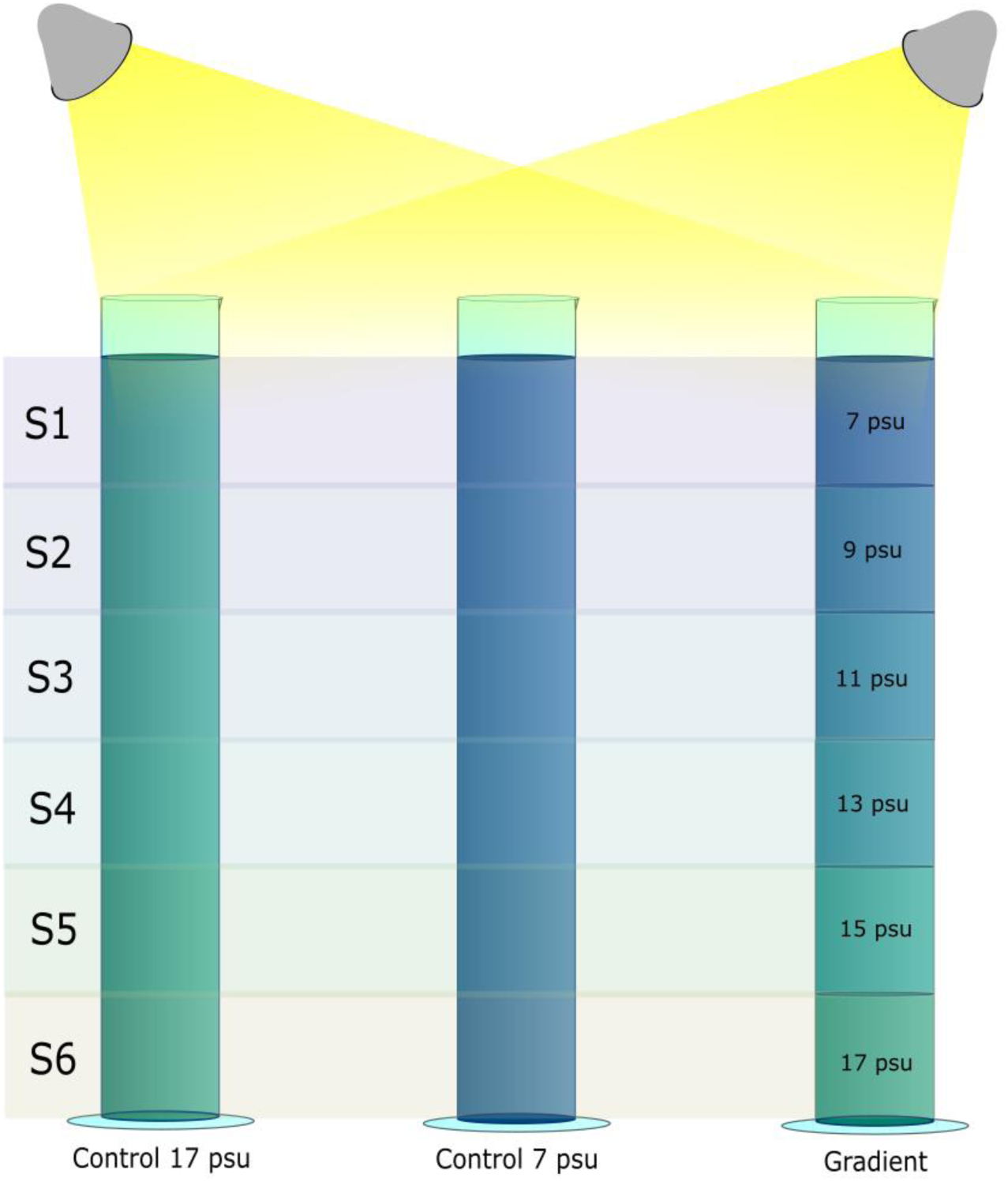
Illustration of the experimental set-up for the behavioural assessment. The sections delimit the different salinity sections in the gradient cylinder, and these delimitations were used to mark the sections in the control cylinders as well.

We selected sixty actively moving larvae belonging to the same batch (i.e., egg collection day) from the incubator and distributed randomly to each of the three cylinders (20 larvae in each) for the behavioural trials. Since cod is not sensitive to red light (Valen et al., 2014), two 16cm long red-light LED lamps were placed in the experimental room to allow the observer to see the larvae in the dark, and two table lamps (52cm tall, 220 lumens) were placed 15cm and at 45° angle above the cylinders illuminating them from the top during the trials in light (Figure 1). The first trial was performed in darkness after 20 minutes of acclimatization period, and the position of each individual larva was marked (under red light) on the wall of the cylinder four times at five-minute intervals. Thereafter, a second trial was done in light conditions after 20 minutes of the acclimatization period using the same larvae. This set of trials (one first in darkness and the second one in light) was repeated three times on the same day, each time using a new set of 60 larvae from the same incubator and belonging to the same batch. We changed the position of the cylinders between replicates to randomize the positions of the light sources in relation to the cylinders. The selected salinity was calculated as the mean position of the 20 larvae in each trial.

### Neutral buoyancy assessment

To assess larval neutral buoyancy, we used three glass columns (86 x 5.5 cm) to create continuous salinity/density gradient columns following the method modified from Coombs (1981). In short, the column was filled up from the bottom, first with low saline water (5 psu) coming from a beaker and pumped by gravity (35-47ml x min-1). At the same time, water in this beaker was stirred with higher saline water (30 psu) coming from an adjacent beaker. The salinity of the water was hence gradually increased in this mixing beaker and pumped into the columns, creating salinity gradient columns from below. The waters were prepared beforehand with artificial seawater and left in the experimental room to attain a temperature of 7°C and to evacuate any gas supersaturation, otherwise forming bubbles in the salinity gradient. We assessed the gradient in each column using five glass beads (different beads in each column), ranging from 7.7 psu to 21,3 psu in buoyancy. Their position was measured to the nearest 2mm on the wall of the column. We estimated the salinity gradient in the column afterwards by interpolating the position of the glass beads against the salinity at which they were neutrally buoyant. Their linearity was high (r^2^ > 0.99) for all measurements. We selected forty-five actively moving larvae from the same incubators and batches used for behavioural assessment. They were sedated in a 0.05 mg benzocaine ml^-1^ water solution and introduced into each column (15 per column) from the top (i.e. from the lowest salinity) and allowed to sink for 10 minutes to the salinity where they were neutrally buoyant. Their position was then noted and related to the salinity derived from the glass beads from its respective column.

To obtain larval body size, we photographed from the side under a microscope (NIKON SMZ800NI) a subsample of 15 larvae from each batch. Standard body length (from the upper lip to the end of the cord) was measured to the nearest 0.01 mm using the image software IC Measure.

### Statistical analyses

We tested whether larval distribution within the cylinders was influenced by salinity treatment (7 psu, 17 psu, or a salinity gradient) and whether light affected this distribution. The mean vertical position of larvae in the cylinders was calculated from four observations under dark conditions (the first trial) and four observations under light conditions (the second trial). This procedure was repeated for each cylinder in every replicate within each batch. We ran a linear mixed model (LMM) with mean section position of the larvae as dependent variable, and cylinder, light and batch as well as their interactions were included as fixed factors. Replicates within each batch were incorporated as a random factor. Statistical significance was computed using the Anova function with Type III ANOVA with Satterthwaite’s method Analyses, performed using car package (Fox and Weisberg, 2019) (Fox and Weisberg, 2019) in R (R Core team 24).

Second, we examined whether there was variation in the larval neutral buoyancy among the batches. We estimated the mean neutral buoyancy for each replicate, and ran a linear mixed model (LMM) with mean neutral buoyancy as dependent variable, batch as fixed effect and replicate as a random effect. We also examined whether there was variation in the larval body length among the batches. We calculated the mean larval body length for each batch and applied a linear model (LM), with mean body length as the dependent variable and batch as the fixed effect. We additionally tested larval buoyancy and size relationship with a linear regression. Statistical significance was computed using the Anova function with Type III ANOVA with Satterthwaite’s method Analyses, performed using car package (Fox and Weisberg, 2019) in R (R Core team 24).

Finally, we examined the roles of neutral buoyancy and light on the salinity position in the gradient cylinder. We ran a linear model (LM) using the mean selected salinity in the gradient cylinder for each replicate under light and dark conditions as the independent variable. Neutral buoyancy was included as a covariate, and light and batch were included as fixed factorial effects, along with the interaction between neutral buoyancy and light. Statistical significance was computed using the Anova function with Type III ANOVA with Satterthwaite’s method Analyses, performed using car package (Fox and Weisberg, 2019) in R (R Core team 24).

## Results

### The effect of salinity and light on the distribution of larvae through the cylinders

We found a significant difference (F_2, 50_ = 866.55, p < 0.001) in the position of larvae among the different cylinders (Fig. 2; Table S2). In the control cylinder with 17 psu, larvae were overall closer to the surface, while in the control cylinder with 7 psu, they were mostly found close to the bottom. In the cylinder with the salinity gradient, the majority of the larvae remained in high salinity conditions (mean psu ± SE of 14.6 ± 0.06 (Fig. 2). Additionally, light had a significant negative phototaxis effect (F_1,50_ = 23.80, p < 0.001; Table S2; Fig. 2) in the position of the larvae across cylinders and variation among the batches was significant as well (F_4, 50_ = 28.92, p < 0.001; Table S2; Fig. 2). The only significant interaction was cylinder:batch interaction (F_8, 50_ = 34.35, p < 0.001; Table S2; Fig. 2), mainly caused by larger variation among the batches in the 17 psu control cylinder (Fig. 2).

**Figure 2:**
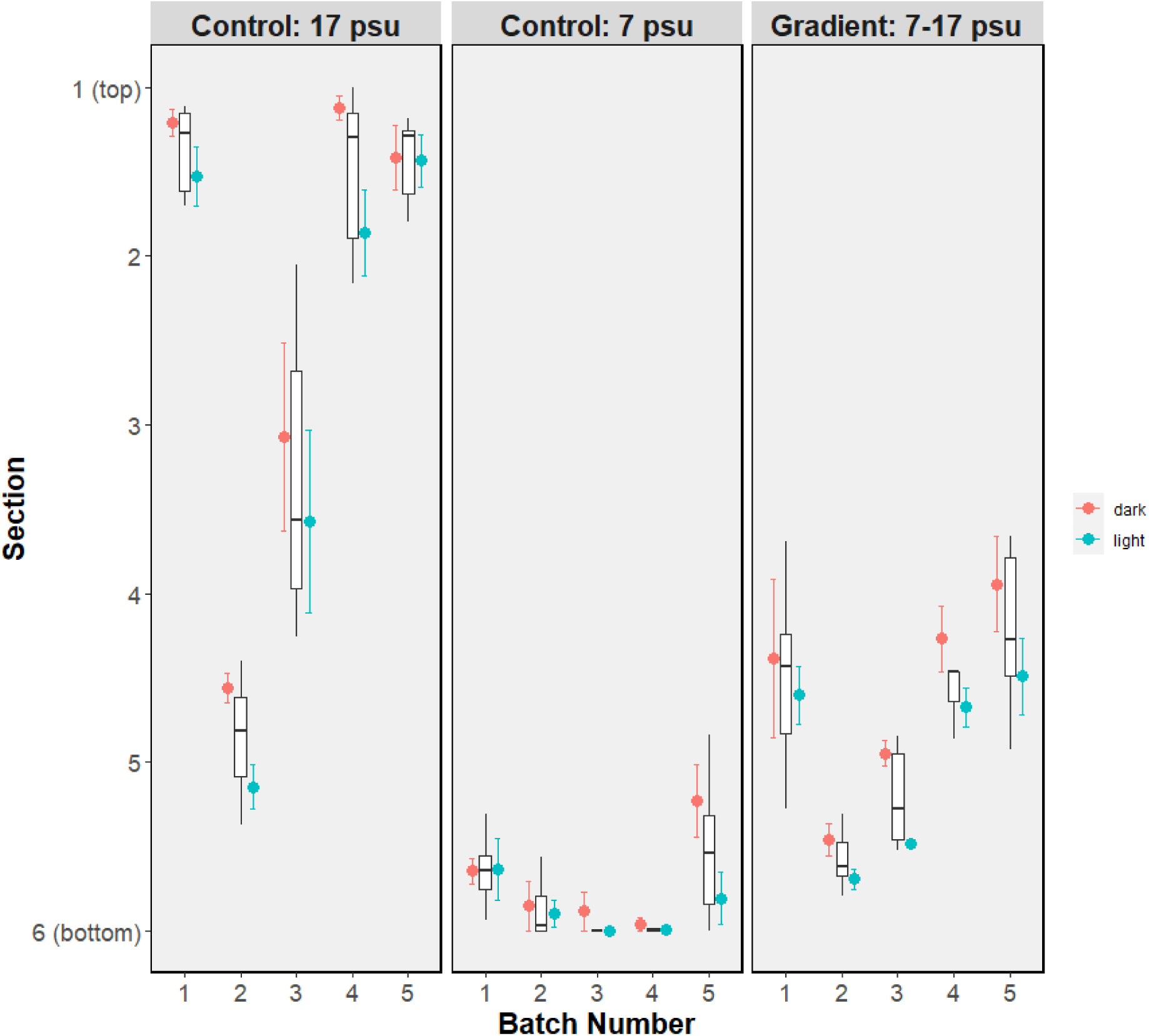
Distribution of larvae across sections in the different cylinders: at 17 psu, at 7 psu and gradient. The mean section position of larvae is shown for each batch and observation condition (red for dark and blue for light condition). Boxplots represent the data distribution within each batch. Boxes illustrate the interquartile ranges, the horizontal line inside the median. The upper and lower whiskers indicate values outside the middle 50%, showing data points within 1.5 times the interquartile range. Dots represent the mean section position of larvae for each batch under light and dark conditions, and the error bars show the standard error of the mean section position.

### Neutral buoyancy and larval size

Mean larval neutral buoyancy was 16.6 ± 0.12 psu, and there was a significant variation among the batches (F_4, 8_ = 29.60, p < 0.001), with mean neutral buoyancy varying between 15 and 18.4 psu among the batches (Fig 3a). Mean larval body length was 4.53 ± 0.03 mm, with significant variation among the batches (F_4, 67_ = 5.50, p < 0.001; Fig. 3b). Neutral buoyancy and larval body length were positively but non-significantly related (y = −24.41 +9.06*x; adjusted R^2^ = 0.6; p = 0.08, n = 5).

**Figure 3:**
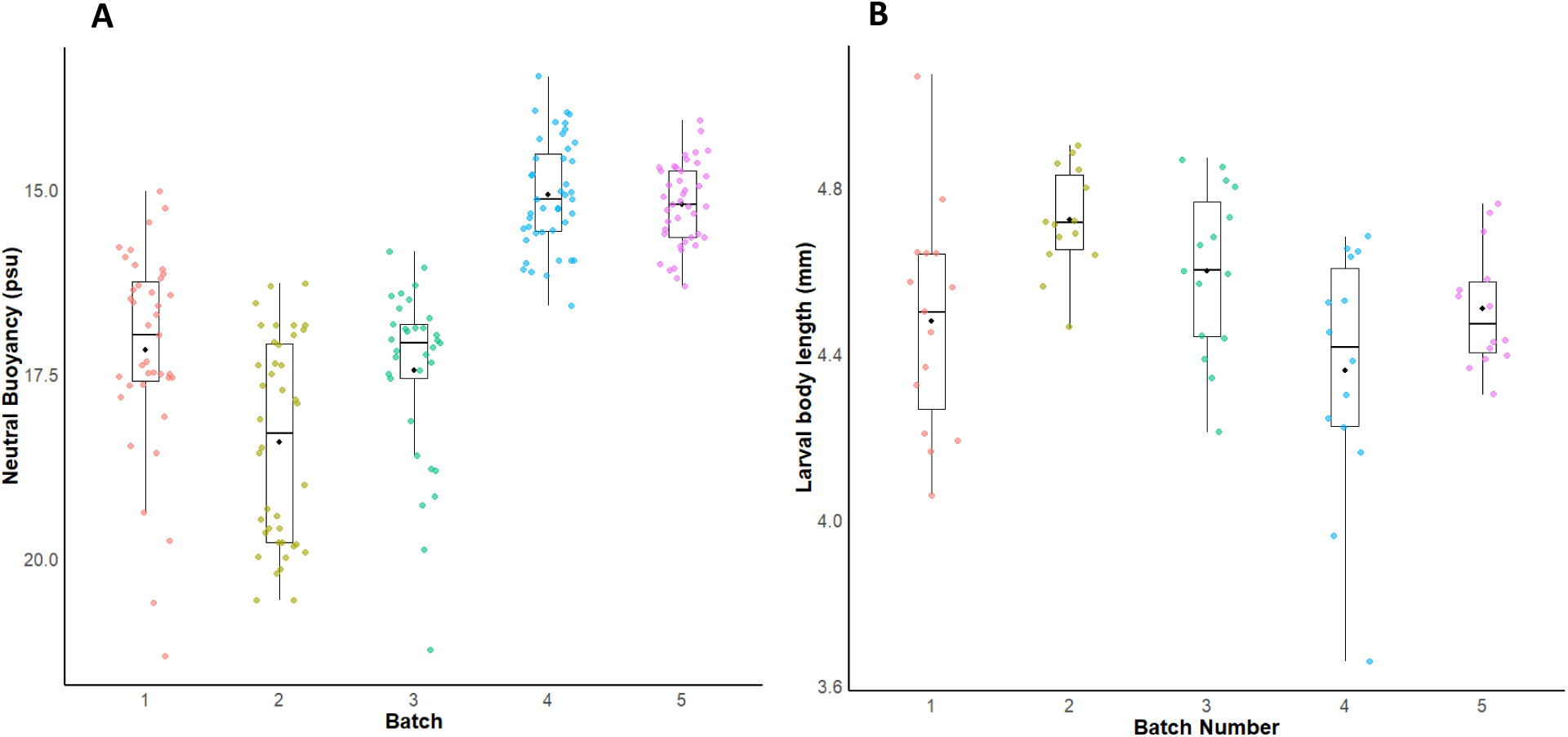
Boxplots of A) neutral buoyancy of larvae (psu) in each batch and B) body length of larvae (mm) in each batch. Boxes inside illustrate the interquartile ranges, the horizontal line inside the median, the upper and lower whiskers indicate values outside the middle 50%, showing data points within 1.5 times the interquartile range. Dots show the data points and the colours the batches.

### Behaviour, neutral buoyancy and light

There was a significant positive effect of neutral buoyancy on the chosen salinity (F_1, 23_ = 48.11, p < 0.001; Table S3; Fig. 4). There was a significant positive effect of light on the chosen salinity (F_1, 23_ = 9.18, p = 0.006). The interaction with neutral buoyancy was not significant (F_1, 23_ = 0.36, p = 0.55; Table S3), indicating that slopes between light and dark conditions remain similar while their intercept differ (Fig. 4). Additionally, significant variation among the batches was detected (F_3, 23_ = 5.46, p = 0.005; Table S3).

**Figure 4:**
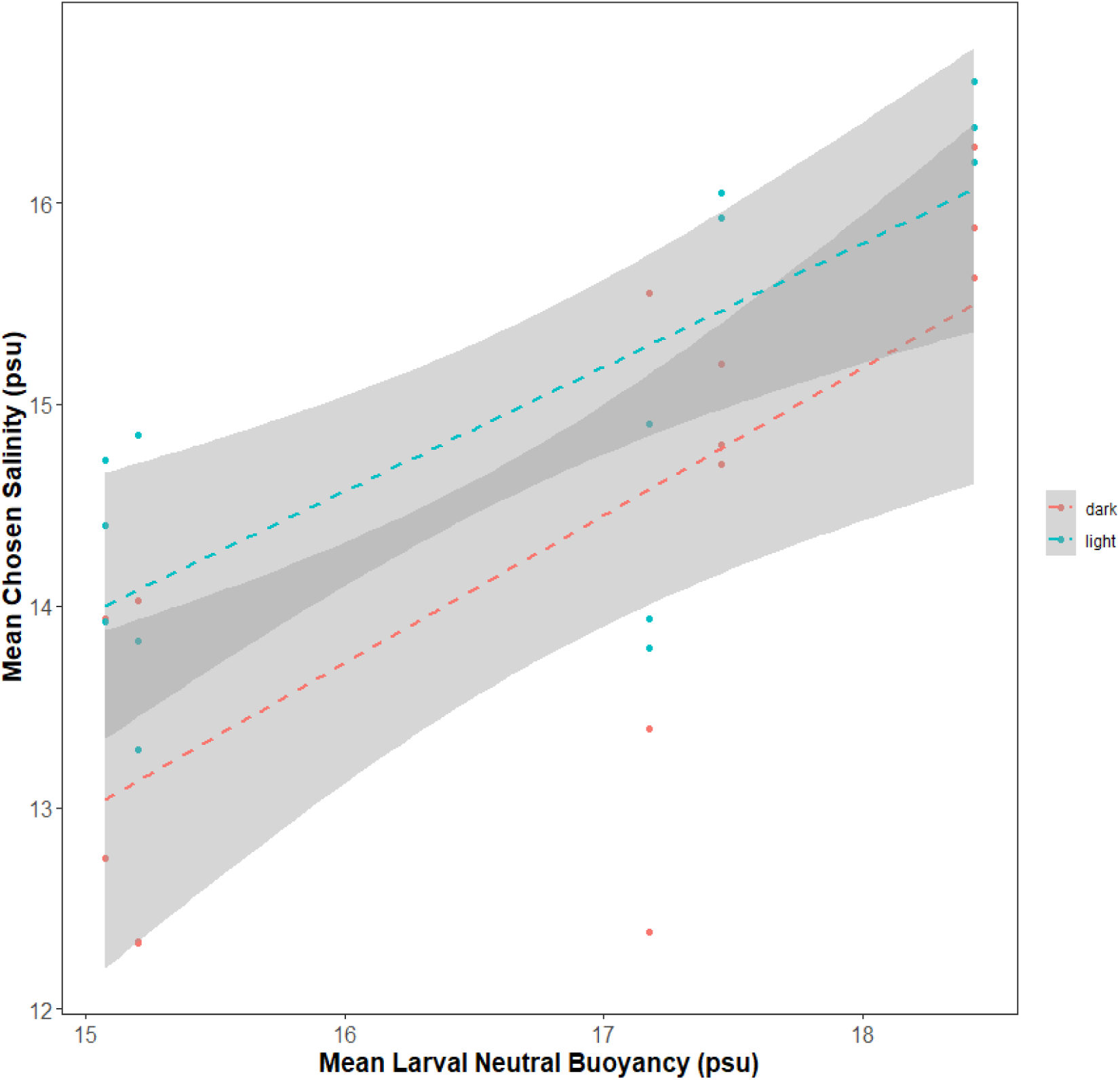
Relationship between the mean neutral buoyancy (psu) and the mean chosen salinity (psu) across batches. The red line represents observations in darkness, while the blue line represents observations in light. Shaded areas indicate 95 % confidence intervals around each regression line.

## Discussion

Our results show a significant variation in the position of larvae between the three cylinders, suggesting that the vertical distribution of the larvae is not random but influenced by environmental salinity and light. We expected larvae to show a preference towards low salinity and a light source. However, the influence of salinity and light was contrary to our expectations. The majority of the larvae in the gradient cylinder were found at salinities of 13-17 psu, with a mean of ∼14 psu, which suggests a preference or choice towards this particular salinity or avoidance of lower salinities. This salinity can be associated with larval neutral buoyancy and reduced osmotic stress. We found that light overall produced negative phototaxis, with larvae moving downwards away from the light source. These results imply that first-feeding larvae remain around the halocline (salinity < 13 psu and dark condition) by the time they start exogenous feeding, rather than migrating upwards for this purpose. Overall, our results suggest that larval neutral buoyancy may play a key role in their vertical distribution and salinity preference, as larvae were primarily observed at salinities close to their neutral buoyancy.

### Larval cod behaviour towards salinity and the influence of light

Our study suggests that first-feeding larvae remain at higher salinity conditions (13 −17 psu), similar to those found below the halocline where cod eggs develop in the Bornholm basin, but lower than the conditions where the larvae were kept before the experiment (17 psu). These results are in line with those reported by Støttrup et al. (2008), showing that first-feeding eastern Baltic cod larvae remained at high salinities. Suitable salinity is crucial for ensuring survival, development and growth at the early life stages of marine fish (Govoni and Forward Jr, 2008; Holliday, 1969; Saillant et al., 2003; Syropoulou et al., 2022), including eastern Baltic cod (Nissling and Westin, 1991b). Optimal salinity for rearing Pacific herring (*Clupea pallasii*) eggs is similar to their body fluid (Holliday, 1969), and in the case of the cod, larval body fluid is similar of the eggs and of the maternal fluid (Davenport et al., 1981). Fish maintain a constant internal osmolarity in their body fluids. In contrast to adults, larval fish lack fully developed and functional gills and excretory organs, and are therefore unable to osmoregulate (Arevalo et al., 2023; Holliday, 1969). Without this ability, they passively gain and lose ions through their skin from the surrounding water (Giffard-Mena et al., 2006; Govoni and Forward Jr, 2008; Varsamos et al., 2005). Avoidance of unsuitable salinity environments by moving towards suitable ones has been observed in adult fish (Herrera et al., 2021; Serrano et al., 2010) and is associated with the avoidance of osmotic stress caused by adverse environmental salinity conditions (Arevalo et al., 2023). First-feeding eastern Baltic cod larvae likely avoid low salinities (e.g., 7 psu) to minimize potential osmotic stress and seek an iso-osmotic environment (12–17 psu) to maintain internal homeostasis. This may explain their tendency to remain in higher-salinity layers, as observed in the gradient cylinder.

Additionally, we did not find evidence that light would enhance movement towards low salinity, an environment rich in food (Grønkjær et al., 1997). Contrary to our initial hypothesis, larvae were detected at approximately 1 psu higher salinity under illuminated conditions in the gradient cylinder. However, similar light avoidance behaviour was observed in the control cylinders. This tendency to avoid light in the control cylinders may indicate that eastern Baltic cod larvae are negatively phototactic, and that such behavioural responses might override the influence of salinity (Burke et al., 1995; Hurst et al., 2009). Atlantic cod larvae rely on light and short distances to detect food (Colton and Hurst, 2010; Ellertsen et al., 1980; Hall et al., 2004), however, a light-avoiding pattern has been observed experimentally in first-feeding Atlantic cod primarily under high light intensities (Ellertsen et al. 1980; Skiftesvik 1994), comparable to those observed in our experiment. Moreover, field studies on eastern Baltic cod showed that the majority of larvae were present below the photic zone and did not show clear larval diel moving patterns (Grønkjær and Wieland, 1997; Huwer et al., 2011).

Larval light avoidance is a common behaviour found in multiple species in field studies (Boehlert and Mundy, 1994; Leis, 1991). It is considered an anti-predator strategy, as their small body size confers high susceptibility to predation, and increased light levels enhance visual detection by predators (Hurst et al., 2009; Saborido-Rey et al., 2003). Phototactic responses at early life stages vary among species (Hurst et al., 2009; Ina et al., 2017; Lough and Potter, 1993; Tielmann et al., 2016), and are further influenced by light intensity, ontogeny, and population. (Burke et al., 1995; Ellertsen et al., 1980; Nicolaisen and Bolla, 2016; Skiftesvik, 1994; Vollset et al., 2009). Atlantic cod larvae shift movements under different light intensities; these responses further varied across developmental stages (Ellertsen et al. 1980; Skiftesvik 1994) and even between populations (Puvanendran, 1999). All these factors play a key role in driving the vertical distribution and movement of larval cod (Ellertsen et al., 1980; Nicolaisen and Bolla, 2016; Skiftesvik, 1994; Vollset et al., 2009). Therefore, our observations are likely specific to the particular light intensity, ontogenetic stage, and population examined. These findings suggest that the role of light in shaping larval behavioural traits is complex and remains poorly understood in eastern Baltic cod and larval fish more broadly, highlighting the need for further investigation.

### Neutral buoyancy and larval size

Mechanisms determining larval neutral buoyancy are not yet fully elucidated, although they are likely influenced by multiple factors such as water content and osmoregulatory capacity regulated through maternal effects (Govoni and Forward Jr, 2008; Saborido-Rey et al., 2003). Newly hatched larvae show similar buoyancy values to the eggs from which they hatched, which is determined during egg maturation in the ovaries and mediated through maternal effect (Govoni and Forward Jr, 2008). Our results indicate that first-feeding stage eastern Baltic cod larvae reach neutral buoyancy at approximately 16.6 psu salinity, consistent with previous findings (Nissling and Vallin, 1996). However, we found that buoyancy was highly variable among the egg batches, varying between 15 and 18.4 psu. High variation in larval neutral buoyancy has been observed in the Atlantic and eastern Baltic cod (Ellertsen et al., 1980; Nissling and Vallin, 1996; Støttrup et al., 2008), as well as in other fish (e.g., Liu H. W. et al., 1993; Tsukamoto K. et al., 2009). For instance, Saborido-Rey et al., (2003) and Helmerson et al. (2023) reported higher values for Atlantic cod (27-35 psu) and western Baltic cod (20-22 psu), respectively, compared to eastern Baltic cod (Nissling et al., 1994; Nissling & Vallin, 1996, Nissling & Westin, 1991a). Differences in buoyancy might reflect the adaptation to the brackish waters of the eastern Baltic cod population (Nissling et al., 1994a). Internal fluid volume and lipid content influence larval buoyancy immediately after hatching (Govoni and Forward Jr, 2008). Internal fluid volume and lipid content are linked to egg size and quality, which vary throughout the spawning season and are directly determined by maternal effects (Pepin et al., 1997; Saborido-Rey et al., 2003). Therefore, variation in parental body size over the extended spawning period suggests that maternal effects related to these traits may account for at least part of the observed variation in larval neutral buoyancy.

Neutral buoyancy is influenced not only by maternal effects but also by larval development and size (Govoni and Forward Jr, 2008), contributing to variation among batches in eastern Baltic cod (Nissling A. et al., 1998; Nissling et al., 1994b). Our results indicate that first-feeding larvae measured approximately 4.5 mm in length, consistent with previous observations in both Atlantic cod (Ellertsen et al., 1980; Fossum, 1986; Hall et al., 2004) and eastern Baltic cod (Grønkjær and Wieland, 1997; Huwer, 2009). Additionally, larval size exhibited variability across different batches and a tendency of larger larvae to exhibit lower buoyancy, confirming the patterns found by (Saborido-Rey et al. (2003) and Vallin & Nissling (2000). As larvae grow, they undergo morphological changes, including alterations in shape and surface area-to-volume ratio, along with increases in structural carbohydrates (chondromucin), bone, calcified cartilage, and muscle, which collectively reduce buoyancy (Govoni and Forward Jr, 2008). Multiple factors, including size, influence and determine larval neutral buoyancy and account for its variation, but more research is needed to estimate how specifically these factors contribute to larval neutral buoyancy.

### Neutral buoyancy and behaviour

Optimal salinity for eastern Baltic cod egg and larval development is > 11 psu (Nissling and Westin, 1991b) and within the range of their neutral buoyancy at 12-17 psu (Nissling and Westin, 1997). Interestingly, our results indicate that larval behaviour is highly influenced by their neutral buoyancy, since larvae remain at salinities similar to their neutral buoyancy. When larval fish reach neutral buoyancy, the osmolarity of their internal fluids approximates that of the surrounding water (Govoni and Forward Jr, 2008). Due to the lack of osmoregulatory ability at this stage, larvae maintain osmotic balance and prevent osmotic stress by actively remaining within salinity ranges close to their internal osmolarity, thereby minimizing exposure to lower salinity environments. In our experiment, larvae were predominantly found near the bottom of the control cylinder under low-salinity conditions (7 psu). In contrast, at higher salinity (17 psu), larvae displayed a wider vertical distribution, frequently occurring near the water surface. This sinking behaviour at lower salinities is consistent with the reduced vertical activity reported by Nissling et al. (1994) linked to an osmotic stress. Larval aggregation in the surface of the cylinder at 17 psu might be a result of their positive buoyancy in this condition, since their neutral buoyancy was around this salinity. Aggregation of larval cod at the surface at salinities similar to their neutral buoyancy has been reported earlier (Nissling et al., 1994b). However, larvae in the gradient cylinder were mainly observed at lower salinities (13–17 psu) compared to those corresponding to their neutral buoyancy range (15 and 18.4 psu), a pattern also observed by Støttrup et al. (2008).

Movement in water is energetically demanding due to its viscosity, with small organisms such as cod larvae incurring disproportionately higher costs compared to larger ones (Facey et al., 2022b; Hunter, 1981; Leis, 2006; Purcell, 1977). The preference for salinities approximating neutral buoyancy may serve to reduce the energetic costs of maintaining position in the water column (Leis, 1991; Saborido-Rey et al., 2003). By remaining at salinities close to their neutral buoyancy, larvae can maintain their position in the water column with little or without swimming activity. Also, these conditions are iso-osmotic, representing optimal salinity for their development and survival (Holliday, 1969). A further possible explanation for the observed larval vertical distribution across cylinders is that the larvae had not yet started active feeding and were not motivated to search for food. However, feeding behaviour in the absence of food has been reported at this developmental stage (Ellertsen et al., 1980).

In summary, our study demonstrates that first-feeding eastern Baltic cod larvae do not exhibit a preference for low salinity or light. Instead, they consistently avoid low salinities even in the presence of light, remaining within the 13–17 psu range, which corresponds closely to their neutral buoyancy. Importantly, this study advances our understanding of the behaviour of eastern Baltic cod during the critical early life stages of this threatened subspecies. Further research, combining laboratory and field studies, in behavioural and physiological responses to low salinity during ontogeny (Bell and Brown, 1995; Ellertsen et al., 1980; Ellien et al., 2011; Rao, 2003; Rønnestad et al., 2013), as well as on feeding behaviour (Bochdansky et al., 2008; Jungbluth et al., 2021), would help increase our knowledge on larval eastern Baltic cod behaviour. Such studies would also provide insight into the environmental conditions most suitable for early life development.

## Acknowledgements

We thank Sebastian Nikitas Politis for input on the experimental design, Göran Anqvist for guidance on the statistical analyses, Gunilla Rosenqvist for providing help with the facilities and Baltic Waters Foundation for financial and logistic support. Last but not least, we thank the staff working on the ReCod project for taking care of the fish and the facilities before and during the experiment, and for their help for making this experiment possible.

