## Supplementary material for "Eastern Baltic cod larvae in a salinity gradient: choice of salinity and the role of neutral buoyancy": Suplemental Table 1, Table 2and Table 3

[3]Present address: Department of Ecology, Environment and Botany, Stockholm University, 114 18 Stockholm, Sweden

### Supplementary material

**Table 1**: Overview of broodstock fish used in this experiment, measured at the end of February and beginning of March 2023 (i.e. before the beginning of the experiment). *n* = number of fish, BL = total length, BW= body weight.

| **Tank** | **BL (cm) (min-max, mean ± SD)** | **BW (g) (min-max, mean ± SD** |
|---|---|---|
| Tank 1 (n = 39) | 57-78, 67.51 ± 5.07 | 2666-6674, 4320.05 ± 1008.04 |
| Tank 2 (n = 53) | 46-63, 54.23 ± 4,17 | 1268-4006, 2231.53 ± 600,63 |

**Table 2**: Results of the Linear Mixed-Effects Model Analysis testing the effect on the position of the larvae of: cylinder (7 psu, 17 psu or gradient), light, batch and replicate (random factor). The table shows the denominator degrees of freedom (NumDF and denDF), F-statistics and P-values for the main effect and interactions, marked with * if significant.

| | **NumDF** | **DenDF** | **F value** | **PR (>F)** | |
|---|---|---|---|---|---|
| Cylinder | 2 | 50 | 866.55 | < 0.001 | *** |
| Batch | 4 | 10 | 28.92 | < 0.001 | *** |
| Light | 1 | 50 | 23.80 | < 0.001 | *** |
| Cylinder:batch | 8 | 50 | 34.35 | < 0.001 | *** |
| Cylinder:light | 2 | 50 | 1.71 | 0.191 | |
| Batch:light | 4 | 50 | 0.40 | 0.811 | |
| Cylinder:batch:light | 8 | 50 | 1.02 | 0.435 | |

**Table 3**. Results of the Linear Model Analysis testing the effect on the mean chosen salinity in the gradient cylinder of: the mean larval neutral buoyancy, light and batch and replicate. The table shows the denominator degrees of freedom (NumDF and denDF), F-statistics and P-values for the main effect and interactions, marked with * if significant.

| | **NumDF** | **DenDF** | **F value** | **Pr (>F)** | |
|---|---|---|---|---|---|
| Mean neutral buoyancy | 1 | 23 | 48.11 | < 0.001 | *** |
| Light | 1 | 23 | 9.18 | 0.006 | ** |
| Batch | 3 | 23 | 5.46 | 0.005 | * |
| Mean neutral buoyancy:light | 1 | 23 | 0.36 | 0.554 | |
